# Improved contiguity of the threespine stickleback genome using long-read sequencing

**DOI:** 10.1101/2020.06.30.170787

**Authors:** Shivangi Nath, Daniel E. Shaw, Michael A. White

**Author notes:** **Corresponding author:** Michael A. White, Department of Genetics, University of Georgia, 120 Green St., Athens, GA 30602.

## Abstract

While the cost and time for assembling a genome have drastically reduced, it still remains a challenge to assemble a highly contiguous genome. These challenges are rapidly being overcome by the integration of long-read sequencing technologies. Here, we use long sequencing reads to improve the contiguity of the threespine stickleback fish *(Gasterosteus aculeatus)* genome, a prominent genetic model species. Using Pacific Biosciences sequencing, we were able to fill over 76% of the gaps in the genome, improving contiguity over five-fold. Our approach was highly accurate, validated by 10X Genomics long-distance linked-reads. In addition to closing a majority of gaps, we were able to assemble segments of telomeres and centromeres throughout the genome. This highlights the power of using long sequencing reads to assemble highly repetitive and difficult to assemble regions of genomes. This latest genome build has been released through a newly designed community genome browser that aims to consolidate the growing number of genomics datasets available for the threespine stickleback fish.

## Introduction

Reference genome assemblies have been invaluable in the discovery of genes, the annotation of regulatory regions, and for providing a scaffold for understanding genetic variation within a species. With the advent of new sequencing technologies and the reduction of cost, there has been a rapid increase in the total number of reference genomes available across taxa. Although it has become much simpler to produce a draft genome assembly, the completion of a high-quality, contiguous assembly remains a great challenge. There are many regions within individual genomes that are unassembled. These regions are enriched for highly repetitive sequence that cannot be assembled using sequencing technologies that produce short fragments (Nagarajan and Pop 2013; Gnerre et al. 2011). Even the most highly refined genomes, like the human genome till have many gaps, which often are composed of long segmental duplications (Schneider et al. 2017).

Long reads from third-generation sequencing technologies have shown promise in spanning highly repetitive regions of genomes, bridging previously intractable gaps in assemblies to improve overall contiguity. Within the human genome, many highly repetitive regions have been resolved, such as pericentromeres (Vollger et al. 2020), complete centromeres (Jain et al. 2018b), telomeres (Jain et al. 2018a) and the entire major histocompatibility complex (Jain et al. 2018a). *De novo* assemblies of highly repetitive Y chromosomes have also become feasible using long-read sequencing (Mahajan et al. 2018; Peichel et al. 2019). Overall, long-read sequencing has enabled telomere-to-telomere chromosome assemblies in multiple species (Liu et al. 2020; Miga et al. 2019). It is clear that hybrid assembly approaches incorporating long-read sequencing have greatly improved contiguity of genomes.

Here we use long-read sequencing to improve the contiguity of the threespine stickleback fish *(Gasterosteus aculeatus)* genome. The threespine stickleback fish has been an important model system to understand evolution, ecology, physiology and toxicology (Wootton 1976; Bell and Foster 1994). The identification of the genetic mechanisms underlying many adaptative traits was facilitated by the release of a high-quality reference genome assembly (Jones et al. 2012). This genome assembly was constructed using paired-end Sanger sequencing reads from multiple genomic libraries. Contigs were scaffolded to genetic linkage maps, which resulted in 21 chromosome-level scaffolds (400.4 Mb), with 60.7 Mb of unplaced scaffolds. The assembly has undergone several revisions, using high-density genetic linkage maps (Roesti et al. 2013; Glazer et al. 2015), and a Hi-C proximity-guided assembly (Peichel et al. 2017). Despite multiple revisions, the latest version of the assembly (v. 4) still contains 13,538 gaps and 20.6 Mb of unplaced scaffolds (Peichel et al. 2017). The gaps between contigs in the chromosome scaffolds likely represent repetitive regions or GC-rich regions, which have been shown to be recalcitrant to traditional assembly methods (Benjamini and Speed 2012; Ross et al. 2013). In order to improve the assembly, we used long-read sequencing to fill gaps in the threespine stickleback genome assembly. We were able to close 76.7% of the gaps, incorporating 13.5% of the previously unplaced scaffolds. Closed gaps were highly accurate, verified through longdistance linked-read information. In addition, we were able to extend sequence of many of chromosomes into telomeres. This assembly represents a noteworthy improvement, allowing researchers to interrogate many previously inaccessible repetitive regions, and highlights the power of long-read sequencing to substantially improve genome contiguity.

## Methods

### Ethics statement

All procedures using threespine stickleback fish were approved by the University of Georgia Animal Care and Use Committee (protocol A2018 10-003-Y2-A5).

### Closing gaps in the reference assembly

Version four of the threespine stickleback reference genome assembly contains 1263 unplaced contigs (chr. Un) that were narrowed to chromosomes but were not placed into specific gaps (there was a total of 3378 chr. Un contigs: 1263 contigs were narrowed broadly to chromosomes and 2115 could not be localized to any chromosome). We used recently available high-coverage long-read sequencing in combination with the 1263 chr. Un contigs that were previously narrowed to chromosomes to fill the remaining gaps in the reference assembly. A male Paxton Lake benthic threespine stickleback fish (Texada Island, British Columbia) was sequenced to approximately 75x coverage (NCBI BioProject database accession PRJNA591630) and assembled into contigs using Canu (Koren et al. 2017). This assembly had a total of 3593 contigs (N50: 683 kb) from across the genome. We closed gaps in the v. 4 threespine stickleback reference assembly using these contigs and LR_Gapcloser with the parameter -a 1 (Xu et al. 2019). We increased the allowed deviation between gap length and the inserted sequence length in order to provide additional flexibility for gap size that was not inferred accurately in the v. 4 genome assembly. LR_Gapcloser fills existing gaps in the genome assembly by identifying contigs which span a gap completely or partially from either end. Three Canu contigs caused a reduction in total chromosome size after placement into gaps. These contigs were likely misassembled by Canu, causing complications in the gap closing. We omitted these three contigs from further analysis. We used BLAT (v. 3.5; Kent 2002) to identify which of the 1263 previously narrowed chr. Un contigs were placed within a gap. We filtered for stringent alignments by only retaining matches where at least 90% of the query length aligned to the assembly and the total aligned region had 2% or less mismatches.

Many chr. Un contigs that were not placed in the v. 4 genome assembly may be represented in the v. 5 assembly if they were contained completely within a PacBio Canu contig (Peichel et al. 2019). To test this, we used BLAT to align the 3378 chr. Un contigs to the new v. 5 assembly. We filtered for stringent alignments by only retaining matches where at least 90% of the query length aligned to the assembly and the total aligned region had 2% or less mismatches. Chr. Un contigs that did not align to the assembly were retained as unassembled and concatenated into a single fasta sequence, with each contig separated by 100 base pairs.

### Validation of the genome assembly using long-distance linked-reads

We verified the revised genome assembly using 10X Genomics long-distance linked-read sequencing from a single female freshwater threespine stickleback fish (Lake Washington, Washington, USA). We extracted high molecular weight DNA from blood using alkaline lysis. Blood was collected from euthanized fish into 0.85x SSC buffer. The cells were collected by centrifuging for two minutes at 2000 xg. Pelleted cells were resuspended in five ml of 0.85x SSC and 27 μl of 20 μg/ml Proteinase K solution. To lyse the cells, five ml of 2x SDS buffer (80mM EDTA, 100mM Tris pH 8.0, and 1% SDS) was added to the suspension and the solution was incubated at 55°C for two minutes. After incubation, 10 ml of buffered phenol/chloroform/isoamyl-alcohol was added to the suspension. The suspension was incubated at room temperature under slow rotation for 30 minutes. The suspension was centrifuged for one minute at 2000 xg at 4°C to separate phases. The aqueous phase was extracted, 10 ml of chloroform was added, and the suspension was rotated for one hour. The chloroform extraction step was repeated twice. After all extractions, the aqueous phase was separated and mixed with ice cold 100% ethanol and one ml of 3M sodium-acetate (pH 5.5). The overall alignment rate of linked-reads to the assembly was 84.4%, resulting in a genome-wide mean read depth of 26.1X.

### Assessing the completeness of the genome assembly

In order to assess the completeness of the genome, we identified universal single copy orthologs (BUSCO) throughout the new assembly, compared to the previous v. 4 assembly (Peichel et al. 2017). BUSCO (v. 3.0.2) was run using default parameters, comparing against the Actinopterygii lineage dataset (4584 total single copy orthologs; OrthoDB v. 9) (Simão et al. 2015). Actinopterygii was used because threespine stickleback fish are teleosts, which is the largest infraclass of Actinopterygii.

### Identification of telomeric sequences

PacBio long reads with highly repetitive regions are often not assembled into contigs. In order to identify telomeric reads, we searched for the ancestral metazoan telomeric motif ‘TTAGGG’ or ‘CCCTAA’ (Traut et al. 2007; Meyne et al. 1989; Moyzis et al. 1988) in the raw PacBio reads. We searched for the motif and their respective counts in each read using the awk command-line utility. Reads were considered for further analyses if they had more than 50 occurrences of the motif. These reads were aligned to the new genome assembly using minimap2 (v. 2.17) (Li 2018) with default parameters to map to PacBio genomic reads (-ax map-pb). Only the primary alignments were retained. Telomeric reads were assigned to a specific chromosome if greater than 10 kb of unique sequence overlapped with one end of a chromosome. Positive telomeric alignments were merged with the new genome assembly. Repetitive sequence content within telomeres was visualized using the dotplot function in Geneious Prime (v2019 1.1) (https://www.geneious.com).

### Identification of centromeric sequences

BLAST+ (blastn; v. 2.7.1) (Camacho et al. 2009) was used to identify the 186 bp threespine stickleback CENP-A monomer repeat (Cech and Peichel 2015) in the PacBio Canu assembled contigs. Contigs containing CENP-A repeats were mapped to the new v. 5 repeat masked assembly (see Genome annotation and repeat masking) using minimap2 (Li 2018) with default parameters to map to PacBio genomic reads (-ax map-pb). Contigs were only retained if greater than 10 kb of sequence mapped uniquely to one chromosome end. The number of CENP-A repeats per chromosome were counted using blastn. Dotplots were generated using Geneious Prime (v2019 1.1) (https://www.geneious.com).

### Genome annotation

Genome features were lifted over from the previous assembly (v. 4) using a hybrid approach. Genome features were first lifted over to the new assembly using the software package flo (Pracana et al. 2017). Most of the features were lifted over successfully (98.1%). We used BLAT to lift over the remaining fraction. The sequence for the features not lifted over with flo were extracted from the version four assembly using samtools faidx. These sequences were then aligned to the new assembly using BLAT. For each feature, the longest alignment was chosen.

Many of the closed gaps were not represented in the previous genome assembly (v. 4) and were therefore unannotated. We annotated these regions using the MAKER (v. 3.01.02) genome annotation pipeline (Cantarel et al. 2008; Holt and Yandell 2011). These annotations combined evidence from multiple RNA-seq transcriptomes, all predicted Ensembl protein sequences (release 95), and *ab inito* gene predictions from SNAP (v. 2006-07-28) (Korf 2004) and Augustus (v. 3.3.2) (Stanke et al. 2006). MAKER was run over three rounds using the RNA-seq transcriptomes and methods previously described (Peichel et al. 2019).

Repeats were annotated across the genome using a combination of RepeatModeler (v. 1.0.11) and RepeatMasker (v. 4.0.5) (http://www.repeatmasker.org). Repeats were first modeled using the default parameters of RepeatModeler. Repeats were then annotated and masked using RepeatMasker with default parameters and the custom RepeatModeler database.

We tested for enrichment of repeats and genes in closed gaps throughout the genome by comparing to randomly drawn 10 Mb segments (we placed 9.9 Mb of sequence within gaps; see Results). We also tested for enrichment of repeats and transposable elements in the remainder of the unplaced chr. Un contigs by comparing to randomly drawn 20 Mb segments throughout the assembled genome (19.6 Mb of chr. Un contigs remained unplaced; see Results). We generated a null distributions by randomly drawing 10,000 segments throughout the genome using bedtools (v. 2.29.2) shuffle (Quinlan and Hall 2010). We then used bedtools intersect to count the number of repeats (with option -c for both 10Mb and 15Mb segments) as well as the number of bases that overlapped genes (with option -wao for 10 Mb segments) within each random segment.

### Data Availability

Long-distance linked-read sequencing and the updated genome assembly are available on the NCBI BioProject database (https://www.ncbi.nlm.nih.gov/bioproject/) under accession number PRJNA639125. The genome assembly is also available for download from the threespine stickleback genome browser (https://stickleback.genetics.uga.edu). All supplemental material has been uploaded to figshare. Reviewer link for accessing the SRA data: https://dataview.ncbi.nlm.nih.gov/object/PRJNA639125?reviewer=agjcut825ub04a0fvdjqj9vpa

## Results and Discussion

### A majority of gaps were closed across the threespine stickleback genome

Version four of the threespine stickleback genome assembly (Peichel et al. 2017) contained 13,538 gaps with an N50 of 91.7kb (total estimated length of gaps: 3,573,762 bp). Using long-read PacBio contigs in conjunction with the 1263 chr. Un contigs that had been narrowed to chromosomes from the previous assembly, we closed 10,394 of the gaps (76.8%), leaving 3,144 gaps in the final assembly (File S1, File S2). In addition to the fully closed gaps, 146 gaps were partially closed. A total of 9,928,283 bases were added to gaps in the assembly. This resulted in an overall greater contiguity of the genome, with a 5.57-fold greater N50 contig length compared to the previous assembly (v. 5 N50: 510.8 kb; v. 4 N50: 91.7 kb) (Table 1).

Genome contiguity and annotation completeness is often assessed by BUSCO (benchmarking universal single copy orthologs) statistics (Simão et al. 2015). We determined if the additional sequence in the new assembly contained coding sequence that improved overall BUSCO metrics. Of the 3640 genes within the database, we found a total of 3521 BUSCO genes in the v.5 assembly. This represented an increase of 99 genes compared to the previous assembly. In addition, the total number of fragmented BUSCO genes decreased to 14, compared to 108 in the previous assembly (Table S1).

Of the 3378 chr. Un contigs from the previous assembly (v. 4), we determined how many were represented in the closed gaps of the new assembly. Of the 3378 contigs, 457 contigs were placed within gaps (13.5%). The previous assembly used a Hi-C-based proximity-guided assembly method that was able to narrow some of the chr. Un contigs (1263) to chromosomes, but was not able to place these contigs into specific gaps (Peichel et al. 2017). We used this information to verify whether our contig placement was corroborated by the Hi-C sequencing. Of the 1263 previously narrowed chr. Un contigs, we placed 90 of into gaps. A majority of these contigs (80.0%) fell within the same chromosome they were assigned to previously by the Hi-C proximity-guided method. This high concordance further confirms our methodology and closure of the gaps.

Across all closed gaps, we added 9.9 Mb of sequence to the genome. 1.0 Mb of this newly added sequence was from chr. Un contigs previously sequenced, but not placed in chromosomes. The remaining 8.9 Mb represented new regions from the long-read sequencing. Many of the gaps in the genome likely represent highly repetitive regions that are challenging to assemble. We compared the repetitive sequence content between the 9.9 Mb of newly added sequence and the remainder of the genome. Indeed, we found newly closed gaps are significantly enriched for repeat sequences (simple and interspersed repeats; 10,000 permutations; *P* < 0.001; Figure S1). Overall, 17.4% of newly added bases contained repetitive DNA compared to 13.5% in the remainder of the genome. Across all newly added gap sequence, we found a total of 1226 protein coding genes. The newly placed regions overall exhibit lower density of coding sequence than the remainder of the genome, although this result was not statistically significant (Figure S1; 10,000 permutations; *P* < 0.083). Only 8.3% of the closed gap bases were contained within coding regions. Across the remainder of the genome 28.3% of bases in the assembly were contained within coding regions. Combined, our results suggest the highly repetitive nature of the sequence contained within these gaps may have prevented assembly of these regions.

Although we closed a majority of gaps in the assembly, we were still unable to determine where 2921 of the chr. Un contigs belonged in the assembly (total length: 19,587,834 bp). One possibility why we were unable to place these contigs is they are even more difficult to assemble due to higher repetitive sequence content. Consistent with this, the unplaced contigs were highly enriched for retrotransposons compared to the placed chr. Un contigs (*P* < 0.001; Figure S2). It is also possible that these contigs represent segments of the genome outside of gaps that are mis-assembled. Our method only focused on closing gaps between contigs. Additional work will be necessary to determine whether these contigs integrate elsewhere in the genome.

### Long-distance linked-reads validated the closing of most gaps

Long-distance linked-reads were used to validate placement of the new sequence. Linked-read molecules that support closure of a gap would exhibit aligned short-reads throughout the closed gap, whereas linked-read molecules that do not support closure of a gap would have aligned short-reads outside of the gap, but a lack of alignment within the gap (Figure 1). The gap closures were highly supported by the linked-read alignments. We only observed 36 gaps (0.3%) that were not supported by linked-reads (i.e. a lack of short-read alignments over the newly added sequence). The remainder of the 10,394 gaps in this analysis that were closed (99.7%) were supported by the long-distance linked-read dataset (Figure 1).

**Figure 1.**
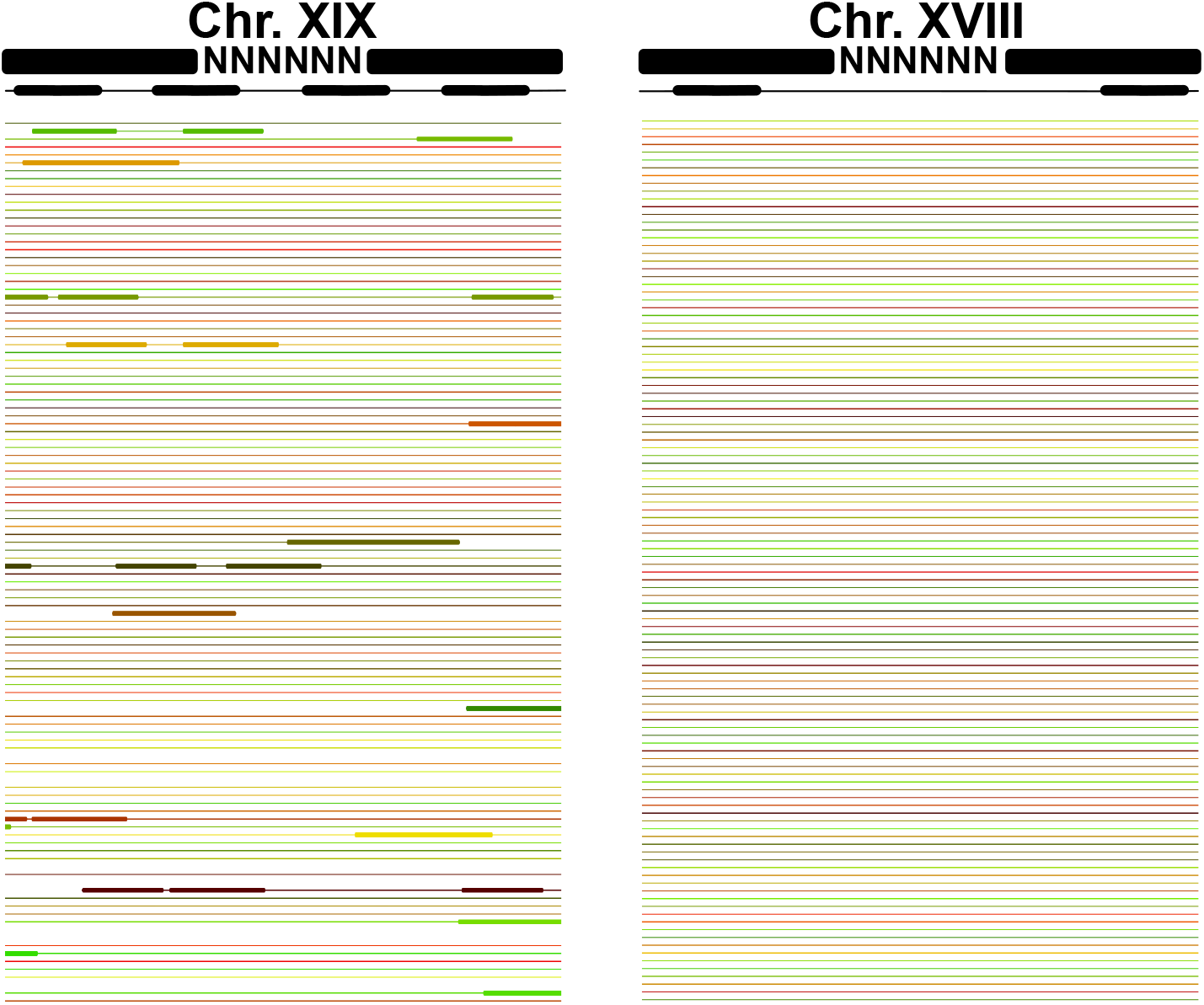
10X Genomics linked-reads validate most of the closed gaps. 99.7% of closed gaps exhibit linked-read alignments throughout the gap region, indicating a correctly closed gap (e.g. Chr. XV: 8,942,107-8,942,560 bp). 0.03% of gaps were not validated by the linked-read sequencing. In these regions, alignments of the linked-reads only occur outside of the gap (e.g. Chr. XII: 20,435,502-20,437,016 bp). A representative schematic outlining how the linked-reads should align is shown in black. The actual aligned linked-reads are shown by bolded color lines. Thin lines indicate gaps between the linked-reads. Average read depth of linked-reads across the genome was 26.IX. A subset of reads aligning is shown here for simplicity.

### Telomere repeats and centromere repeats were identified within PacBio long reads

The telomeres of threespine stickleback fish contain a tandemly repeated G-rich hexanucleotide repeat that is conserved across metazoans (Meyne et al. 1989; Moyzis et al. 1988; Traut et al. 2007; Ocalewicz 2013). Although DNA probes targeting these repeats clearly hybridize at the ends of all chromosomes in threespine stickleback fish, the underlying sequence of these regions is missing from the genome assembly. We therefore searched for the ancestral metazoan telomeric motif ‘TTAGGG’ or ‘CCCTAA’ in the raw PacBio reads to identify putative telomere caps (Konrad et al. 2011; Ocalewicz et al. 2011). We identified 3525 PacBio reads that contained telomere motifs. Seven of these reads contained enough unique sequence to align to the end of individual chromosomes (chromosomes IV, VII, VIII, X, XIV, XV, XVII). These reads showed an abundance of the ancestral metazoan telomeric motif at one end with little to no higher order structure (Figure 2; Figure S3). The telomeric motif was repeated 114-492 times throughout the sequence on different chromosomes.

**Figure 2.**
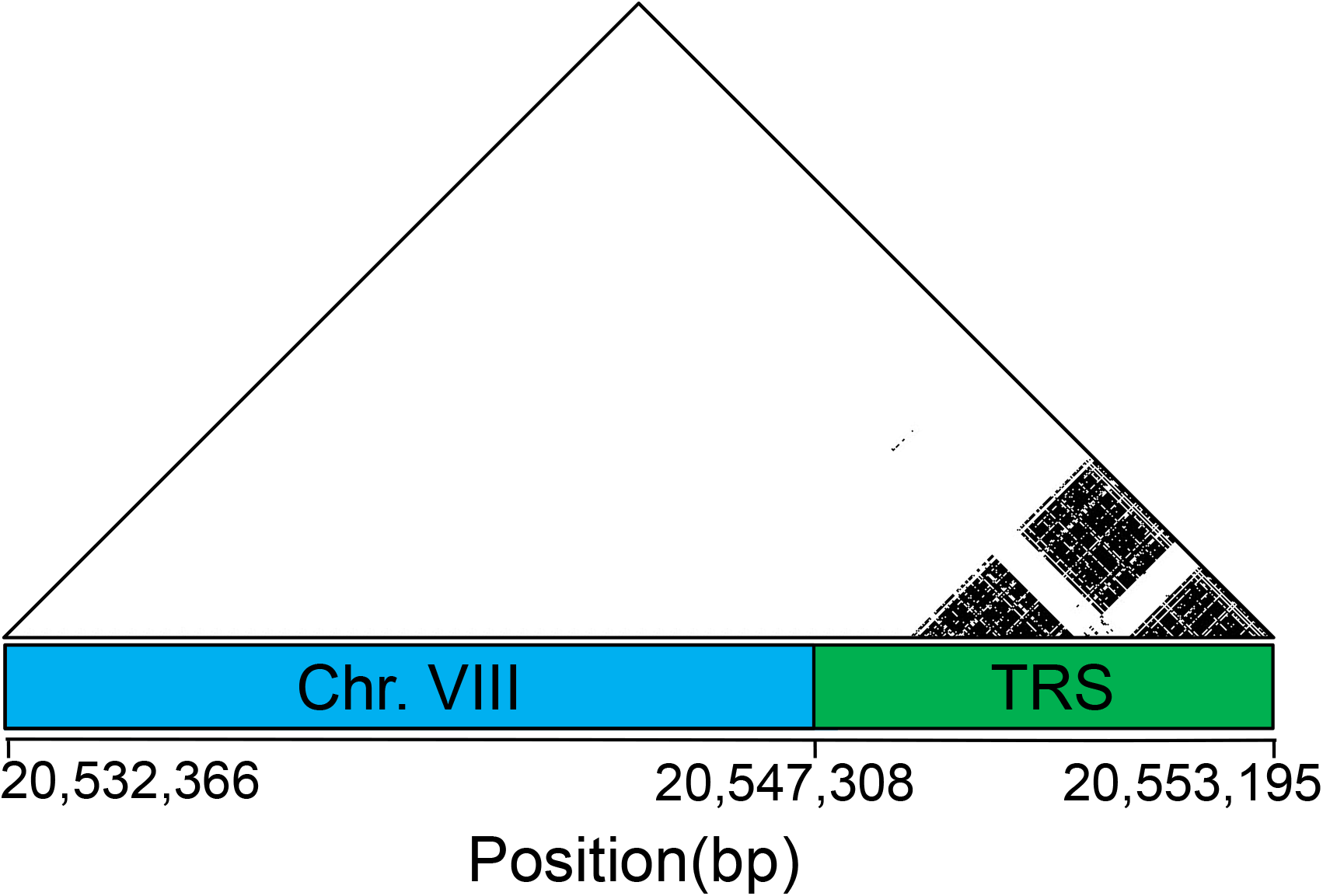
Telomeres exhibit a high density of the conserved metazoan telomere motif. Dots represent 100% sequence identity between matching windows of 15 bp. The blue box represents the end of chr. VII where the long read aligns uniquely to positions 30,722,87630,767,092. The green box denotes a ~10 kb segment rich with telomeric repeat sequence (TRS). The remaining telomeres are shown in Figure S3.

We also searched for centromere repeats within the PacBio assembled contigs. We identified the core 186 bp CENP-A repeat (Cech and Peichel 2015) within 91 contigs (the length of repetitive DNA among contigs ranges from 12.61 - 125.17 kb). 48 of these contigs contained enough unique sequence to align to all 21 chromosomes (Figure 3; File S3; File S4). 11 chromosomes had centromere contigs that map to both sides of the gap, 9 chromosomes had a centromere contig only on one end of the centromere, and one contig contained a full centromere sequence, spanning the entire gap (chromosome IX). Interestingly, on many of the chromosomes, the repeat length was long enough to discern clear higher order repeat structure (Figure 3; Figure S4). Our results are similar to the variability in higher order repeat among the autosomes and X chromosome of humans (Hartley and O’Neill 2019; Willard 1985; Willard et al. 1986; Alexandrov et al. 1993; Shepelev et al. 2015). We detected multiple contigs mapping to either end of the centromere gap for all chromosomes (File S3, File S4), indicating the male fish used for sequencing is likely heterozygous for centromeric arrays. This is consistent with high polymorphism of centromere arrays observed within other species (Willard et al. 1986; Greig et al. 1991; Mahtani and Willard 1990; Devilee et al. 1988; Wevrick and Willard 1989).

**Figure 3.**
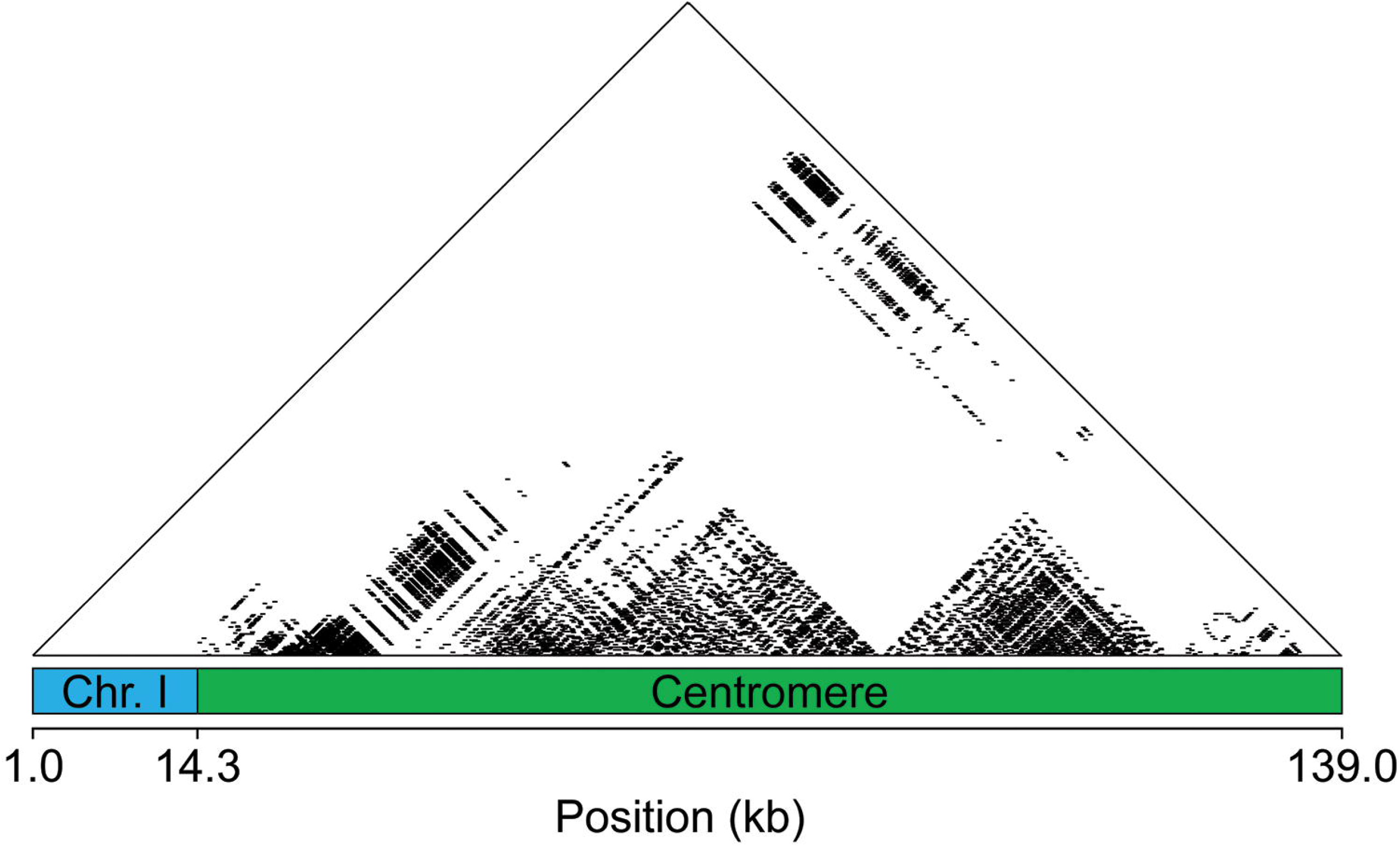
Centromeres display higher order repeat structure. On chromosome I, the centromere contig contains 673 186 bp monomer repeats. Sequence identity between repeats is depicted by black dots matching windows of 300 bp with 100% sequence identity. The blue region denotes the end of chromosome I (20,330,007-20,344,665 bp) with unique sequence. The green region is the newly aligned centromere contig. The remaining centromeres are shown in Figure S4.

Y chromosomes in mammals have also been documented to have highly variable centromeric repeats that are divergent from their counterparts across the remainder of the genome (Wolfe et al. 1985; Pertile et al. 2009; Miga et al. 2014). Assembly of segments of the threespine stickleback Y chromosome centromere (Peichel et al. 2019) revealed an alpha satellite monomer repeat that was divergent from the consensus monomeric repeat identified from the remainder of the genome (Cech and Peichel 2015). With the assembly of larger tracks of centromeric sequence from the autosomes and the X chromosome, we now show the Y chromosome centromere is also divergent from the remainder of the genome at the level of higher order repeats (Peichel et al. 2019), matching other rapidly evolving Y chromosomes. Although our assembly has uncovered a large fraction of the centromeric sequence for each chromosome, we were unable to assemble complete centromere sequences outside of the 46.5 kb centromere of chromosome IX. It therefore remains unknown how centromere length varies throughout the threespine stickleback genome. Complete characterization of the centromeric repetitive arrays will be aided by future sequencing of ultra-long reads (Jain et al. 2018b; Miga et al. 2019).

## Conclusions

By using long-read sequencing we were able to substantially improve the overall contiguity of the threespine stickleback genome, increasing the N50 length of contigs over fivefold. Our assembly also highlights the power of using long-read sequencing technologies to assemble previously inaccessible regions of the genome, like centromeres and telomeres. We have released this assembly through a new threespine stickleback fish community genome browser (https://stickleback.genetics.uga.edu). This resource will be a useful addition to the rapidly expanding functional genomics toolkit available in threespine stickleback fish.

## Acknowledgements

This research was funded by the National Science Foundation IOS 1645170 (M.A.W.), the Office of the Vice President of Research at the University of Georgia (M.A.W.), and the University of Georgia Research Foundation (D.E.S.). We thank the Georgia Genomics and Bioinformatics Core at the University of Georgia for help with the long-distance linked-read sequencing. We also thank Brigitte Hofmeister and the Franklin College Office of Information Technology at the University of Georgia for help building the threespine stickleback genome browser.

## References

Alexandrov, I.A., LI. Medvedev, T.D. Mashkova, L.L. Kisselev, L.Y. Romanova et al., 1993 Definition of a new alpha satellite suprachromosomal family characterized by monomeric organization. Nucleic Acids Res 21 (9):2209–2215.

Bell, M., and S.A. Foster, 1994 The evolutionary biology of the threespine sticklebacks. Oxford University Press.

Benjamini, Y., and T.P. Speed, 2012 Summarizing and correcting the GC content bias in high-throughput sequencing. Nucleic Acids Res 40 (10):e72.

Camacho, C., G. Coulouris, V. Avagyan, N. Ma, J. Papadopoulos et al., 2009 BLAST+: architecture and applications. BMC Bioinformatics 10:421.

Cantarel, B.L., I. Korf, S.M. Robb, G. Parra, E. Ross et al., 2008 MAKER: an easy-to-use annotation pipeline designed for emerging model organism genomes. Genome Res 18 (1):188–196.

Cech, J.N., and C.L. Peichel, 2015 Identification of the centromeric repeat in the threespine stickleback fish (Gasterosteus aculeatus). Chromosome Res 23 (4):767–779.

Devilee, P., T. Kievits, J.S. Waye, P.L. Pearson, and H.F. Willard, 1988 Chromosome-specific alpha satellite DNA: isolation and mapping of a polymorphic alphoid repeat from human chromosome 10. Genomics 3 (l):l–7.

Glazer, A.M., E.E. Killingbeck, T. Mitros, D.S. Rokhsar, and C.T. Miller, 2015 Genome Assembly Improvement and Mapping Convergently Evolved Skeletal Traits in Sticklebacks with Genotyping-by-Sequencing. G3 (Bethesda) 5 (7):1463–1472.

Gnerre, S., I. Maccallum, D. Przybylski, F.J. Ribeiro, J.N. Burton et al., 2011 High-quality draft assemblies of mammalian genomes from massively parallel sequence data. Proc Natl AcadSci USA 108 (4):1513–1518.

Greig, G.M., S. Parikh, J. George, V.E. Powers, and H.F. Willard, 1991 Molecular cytogenetics of alpha satellite DNA from chromosome 12: fluorescence in situ hybridization and description of DNA and array length polymorphisms. Cytogenet Cell Genet 56 (3–4):144–148.

Hartley, G., and R.J. O’Neill, 2019 Centromere Repeats: Hidden Gems of the Genome. Genes (Basel) 10 (3).

Holt, C., and M. Yandell, 2011 MAKER2: an annotation pipeline and genome-database management tool for second-generation genome projects. BMC Bioinformatics 12:491.

Jain, M., S. Koren, K.H. Miga, J. Quick, A.C. Rand et al., 2018a Nanopore sequencing and assembly of a human genome with ultra-long reads. Nat Biotechnol 36 (4):338–345.

Jain, M., H.E. Olsen, D.J. Turner, D. Stoddart, K.V. Bulazel et al., 2018b Linear assembly of a human centromere on the Y chromosome. Nat Biotechnol 36 (4):321–323.

Jones, F.C., M.G. Grabherr, Y.F. Chan, P. Russell, E. Mauceli et al., 2012 The genomic basis of adaptive evolution in threespine sticklebacks. Nature 484 (7392):55–61.

Kent, W.J., 2002 BLAT--the BLAST-like alignment tool. Genome Res 12 (4):656–664.

Konrad, O., W. Piotr, F.-S. Grazyna, and J. Malgorzata, 2011 Chromosomal location of Ag/CMA 3-NORs, 5S rDNA and telomeric repeats in two stickleback species, pp. 12–19. Italian Journal of Zoology.

Koren, S., B.P. Walenz, K. Berlin, J.R. Miller, N.H. Bergman et al., 2017 Canu: scalable and accurate long-read assembly via adaptive. Genome Res 27 (5):722–736.

Korf, I., 2004 Gene finding in novel genomes. BMC Bioinformatics 5:59.

Li, H., 2018 Minimap2: pairwise alignment for nucleotide sequences. Bioinformatics 34 (18):3094–3100.

Liu, J., A.S. Seetharam, K. Chougule, S. Ou, K.W. Swentowsky et al., 2020 Gapless assembly of maize chromosomes using long-read technologies. Genome Biol 21 (1):121.

Mahajan, S., K.H. Wei, M.J. Nalley, L. Gibilisco, and D. Bachtrog, 2018 De novo assembly of a young Drosophila Y chromosome using single-molecule sequencing and chromatin conformation capture. PLoS Biol 16 (7):e2006348.

Mahtani, M.M., and H.F. Willard, 1990 Pulsed-field gel analysis of alpha-satellite DNA at the human X chromosome centromere: high-frequency polymorphisms and array size estimate. Genomics 7 (4):607–613.

Meyne, J., R.L. Ratliff, and R.K. Moyzis, 1989 Conservation of the human telomere sequence (TTAGGG)n among vertebrates. Proc Natl Acad Sci USA 86 (18):7049–7053.

Miga, K.H., S. Koren, A. Rhie, M.R. Vollger, A. Gershman et al., 2019 Telomere-to-telomere assembly of a complete human X chromosome. bioRxiv:135928.

Miga, K.H., Y. Newton, M. Jain, N. Altemose, H.F. Willard et al., 2014 Centromere reference models for human chromosomes X and Y satellite arrays. Genome Res 24 (4):697–707.

Moyzis, R.K., J.M. Buckingham, L.S. Cram, M. Dani, L.L. Deaven et al., 1988 A highly conserved repetitive DNA sequence, (TTAGGG)n, present at the telomeres of human chromosomes. Proc Natl Acad Sci USA 85 (18):6622–6626.

Nagarajan, N., and M. Pop, 2013 Sequence assembly demystified. Nat Rev Genet 14 (3):157–167.

Ocalewicz, K., 2013 Telomeres in fishes. Cytogenet Genome Res 141 (2-3):114–125.

Ocalewicz, K., P. Woznicki, G. Furgala-Selezniow, and M. Jankun, 2011 Chromosomal location of Ag/CMA 3-NORs, 5S rDNA and telomeric repeats in two stickleback species. Italian Journal of Zoology:(12–19).

Peichel, C.L., S.R. McCann, J.A. Ross, A.F.S. Naftaly, J.R. Urton et al., 2019 Assembly of a young vertebrate Y chromosome reveals convergent signatures of sex chromosome evolution. bioRxiv:2019.2012.2012.874701.

Peichel, C.L., S.T. Sullivan, I. Liachko, and M.A. White, 2017 Improvement of the Threespine Stickleback Genome Using a Hi-C-Based Proximity-Guided Assembly. J Hered 108 (6):693–700.

Pertile, M.D., A.N. Graham, K.H. Choo, and P. Kalitsis, 2009 Rapid evolution of mouse Y centromere repeat DNA belies recent sequence stability. Genome Res 19 (12):2202–2213.

Pracana, R., A. Priyam, I. Levantis, R.A. Nichols, and Y. Wurm, 2017 The fire ant social chromosome supergene variant Sb shows low diversity but high divergence from SB. Mol Ecol 26 (11):2864–2879.

Quinlan, A.R., and I.M. Hall, 2010 BEDTools: a flexible suite of utilities for comparing genomic features. Bioinformatics 26 (6):841–842.

Roesti, M., D. Moser, and D. Berner, 2013 Recombination in the threespine stickleback genome--patterns and consequences. Mol Ecol 22 (11):3014–3027.

Ross, M.G., C. Russ, M. Costello, A. Hollinger, N.J. Lennon et al., 2013 Characterizing and measuring bias in sequence data. Genome Biol 14 (5):R51.

Schneider, V.A., T. Graves-Lindsay, K. Howe, N. Bouk, H.C. Chen et al., 2017 Evaluation of GRCh38 and de novo haploid genome assemblies demonstrates the enduring quality of the reference assembly. Genome Res 27 (5):849–864.

Shepelev, V.A., LI. Uralsky, A.A. Alexandrov, Y.B. Yurov, E.l. Rogaev et al., 2015 Annotation of suprachromosomal families reveals uncommon types of alpha satellite organization in pericentromeric regions of hg38 human genome assembly. Genom Data 5:139–146.

Simão, F.A., R.M. Waterhouse, P. Ioannidis, E.V. Kriventseva, and E.M. Zdobnov, 2015 BUSCO: assessing genome assembly and annotation completeness with single-copy orthologs. Bioinformatics 31 (19):3210–3212.

Stanke, M., A. Tzvetkova, and B. Morgenstern, 2006 AUGUSTUS at EGASP: using EST, protein and genomic alignments for improved gene prediction in the human genome. Genome Biol 7 Suppl 1:S11.11–18.

Traut, W., M. Szczepanowski, M. Vítková, C. Opitz, F. Marec et al., 2007 The telomere repeat motif of basal Metazoa. Chromosome Res 15 (3):371–382.

Vollger, M.R., G.A. Logsdon, P.A. Audano, A. Sulovari, D. Porubsky et al., 2020 Improved assembly and variant detection of a haploid human genome using single-molecule, high-fidelity long reads. Ann Hum Genet 84 (2):125–140.

Wevrick, R., and H.F. Willard, 1989 Long-range organization of tandem arrays of alpha satellite DNA at the centromeres of human chromosomes: high-frequency array-length polymorphism and meiotic stability. Proc Natl Acad Sci USA 86 (23):9394–9398.

Willard, H.F., 1985 Chromosome-specific organization of human alpha satellite DNA. Am J Hum Genet 37 (3):524–532.

Willard, H.F., J.S. Waye, M.H. Skolnick, C.E. Schwartz, V.E. Powers et al., 1986 Detection of restriction fragment length polymorphisms at the centromeres of human chromosomes by using chromosome-specific alpha satellite DNA probes: implications for development of centromere-based genetic linkage maps. Proc Natl Acad Sci USA 83 (15):5611–5615.

Wolfe, J., S.M. Darling, R.P. Erickson, I.W. Craig, V.J. Buckle et al., 1985 Isolation and characterization of an alphoid centromeric repeat family from the human Y chromosome. J Mol Biol 182 (4):477–485.

Wootton, R., 1976 The Biology of Sticklebacks. Academic Press.

Xu, G.C., T.J. Xu, R. Zhu, Y. Zhang, S.Q. Li et al., 2019 LR_Gapcloser: a tiling path-based gap closer that uses long reads to complete genome assembly. Gigascience 8 (1).

